# The microtubular preprophase band recruits Myosin XI to the division site for plant cytokinesis

**DOI:** 10.1101/2022.11.08.515512

**Authors:** Calvin Haoyuan Huang, Felicia Lei Peng, Yuh-Ru Julie Lee, Bo Liu

**Affiliations:** Department of Plant Biology, College of Biological Sciences, University of California, Davis, CA 95616, USA; Department of Genetics, Perelman School of Medicine, University of Pennsylvania, Philadelphia, PA 19104, USA

## Abstract

Plant growth is dependent on oriented cell divisions that employ the microtubular preprophase band (PPB) to position the cell plate. It has been intriguing how this transient cytoskeletal array imprints the spatial information to be read by the cytokinetic phragmoplast at later stages of mitotic cell division. In *Arabidopsis thaliana*, we discovered that the PPB recruited the Myosin XI motor MYA1 to the cortical division site where it joined microtubule-associated proteins and motors to form a ring of prominent cytoskeletal assemblies which received the expanding phragmoplast. This regulatory function of MYA1 in phragmoplast guidance is dependent on intact actin filaments. The discovery of these assemblies revealed the mechanism underlying how two dynamic cytoskeletal networks govern PPB-dependent division plane orientation during vegetative growth in flowering plants.

**ONE-SENTENCE SUMMARY:** Myosin XI joins microtubule-associated proteins and motors to form cortical assemblies to demarcate the cell division site.

---

Plant growth is dependent on the production of new cells in physiologically important orientations in order to build tissues in a spatially regulated manner, in part because plant cells which are constrained by rigid cell wall that prevents cell locomotion. From embryogenesis to organogenesis, plant cytokinesis employs a transient microtubule-based cytoskeletal array named the preprophase band (PPB) at the cell cortex. The PPB is established during the G2 phase to demarcate the division site and disassembled concomitantly with nuclear envelope breakdown (NEB) towards the end of prophase (*1*). During vegetative growth, the PPB is employed in both proliferating divisions to produce more identical cells and formative division to produce two daughter cells with distinct fates. The PPB-demarcated cortical division site (CDS) is read by the cytokinetic apparatus phragmoplast so that the cell plate synthesized by the latter will be inserted into plasma membrane at the CDS to partition the two daughter cells. The PPB is therefore of paramount importance for tissue generation and growth throughout the life of a plant (*2*).

Previously, a pharmacological study informed us that the translation of the PPB microtubule array into the CDS after PPB disassembly is dependent on actin microfilaments (F-actin) in plant cells (*3*). F-actin is detected in the PPB at early stages of its development but disappear from the mature PPB and are undetectable at the CDS (*4*). It has been enigmatic what role F-actin plays in allowing the expanding phragmoplast with the developing cell plate to precisely recognize the CDS at later stages of cell division. In the meantime, proteins like the microtubule-associated protein TAN1 and its interacting Kinesin-12 motor POK1 persist at the CDS after PPB disassembly and play critical roles in the maintenance of the CDS established by the PPB (*5*). However, in the context of cell division plane determination, it remains unclear how the function of these microtubule-associated factors is integrated with that of F-actin.

The crosstalk between the two cytoskeletal systems during plant cell division may be mediated by myosin motors as evidenced by the spindle and phragmoplast association of the Myosin VIII and Myosin XI motors in the moss *Physcomitrium patens* and Myosin XI-K in the angiosperm *Arabidopsis thaliana* (*6–8*). In these two plant models, simultaneous losses of multiple Myosin VIII and Myosin XI, respectively, led to weak division plane misalignment phenotypes. A recent study also revealed of the Myosin XI isoform OPAQUE1 plays a critical role in phragmoplast guidance during asymmetric division for stomatal development in maize (*9*). In *A. thaliana*, different Myosin XI isoforms including Myosin XI-K were predicted to acquire specialized functions due to the diversification of their sequences and enzymatic properties (*10*). A published study revealed that F-actin generates the force for the development of the narrow PPB microtubule array from a wide one (*11*). Therefore, we hypothesized that actin-based motors play critical roles in division plane orientation and functionally redundant Myosin XI motors mediated the actin-microtubule interaction at the cell cortex during mitotic cell division in plants. To test this hypothesis, we captured the dynamics of a mitotically active Myosin XI motor and discovered its microtubule-coupled function in cytokinesis. Our findings uncovered a long sought-after mechanism underlying the determination of cell division planes in plant cells.

Four of the 13 Myosin XI motors in *A*. *thaliana, Myosin XI-1/MYA1, Myosin XI-2/MYA2, Myosin XI-I*, and *Myosin XI-K/XI-K* exhibit elevated expression patterns in vegetative tissues. The corresponding quadruple mutant (MyoXI-4KO hereafter) shows significant reduction in growth when compared to wild-type plants or mutants of lower orders (*12, 13*) Because F-actin plays a role in microtubule organization in both the PPB at prophase and the spindle midzone at early stages of cytokinesis (*11, 14*), we tested whether this MyoXI-4KO mutant experienced deficiencies in microtubule-associated activities. To do so, we grew seedlings on medium supplemented with 150 nM oryzalin, a microtubule-disrupting herbicide. The MyoXI-4KO mutant seedlings showed exacerbated growth defects in the presence of oryzalin as evidenced by shorter roots when compared to the wild-type control although mutant roots did not grow significantly differently without the oryzalin challenge (Figure 1A, Supplemental Figure 1). To test whether this oryzalin hypersensitive phenotype was linked to the mutations in the myosin genes, we introduced a construct for MYA1 expression in a GFP (green fluorescent protein) fusion under the control of the *MYA1* promoter into the MyoXI-4KO mutant. *MYA1* was chosen because of its highest expression level in vegetative tissues (*12*). The growth phenotype of the MyoXI-4KO mutant was suppressed by MYA1-GFP expression as the transgenic line, referred hereafter as rescue, was comparable to the wild-type control (Figure 1A, Supplemental Figure 1), confirming that the *mya1* mutation had a causative relationship with oryzalin hypersensitivity.

**Figure 1.**
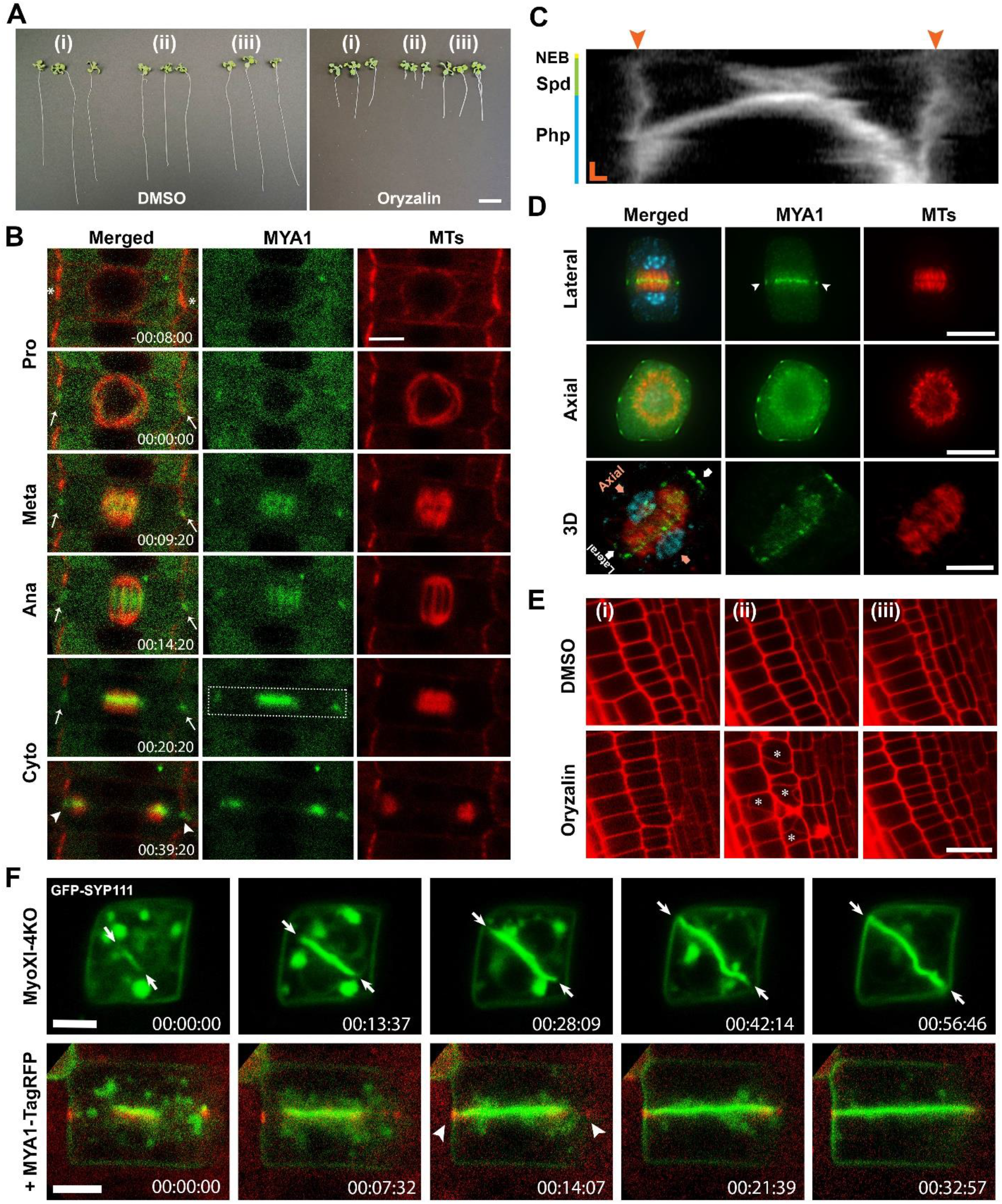
The Myosin XI motor MYA1 decorates the cortical division site and plays a critical role in cell division plane determination in *A*. *thaliana*. Scale bars: 1 cm (A), 5 μm (B, D, F), and 20 μm (E). (A) The Myosin XI quadruple mutant (MyoXI-4KO) displays hypersensitivity to oryzalin. Seedlings of wild-type (i), MyoXI-4KO (ii), and MyoXI-4KO expressing MYA1-GFP (iii) were grown on media without and with 150 nM oryzalin which causes severe inhibition of root growth when compared to the wild-type control and rescue line expressing the MYA1-GFP transgene. Quantitative analysis of root length is shown in Supplemental Figure 1A. (B) MYA1 exhibits a cell cycle-dependent localization pattern. MYA1-GFP localizes to the CDS at late prophase and continuously marks the CDS until the phragmoplast reaches the site. Cell cycle stages are marked as Pro (prophase), Meta (metaphase), Ana (anaphase), and Cyto (cytokinesis). (C) A Kymographic view reveals the MYA1-GFP dynamics in the area outlined by the dotted box in (B). Prior to nuclear envelope breakdown (NEB), MYA1-GFP emerges at the CDS (arrowheads) and continuously associates with it. Following the assembly of the spindle apparatus (Spd), the MYA1 signal arises in the spindle midzone and is later continued by conspicuous decoration of the phragmoplast (Php) midzone. The vertical bar represents 10 min and the horizontal bar marks 1 μm in the Kymogram. (D) Dual immunolocalizations of MYA1-GFP (green) and microtubules (red) in fixed cytokinetic cells with DNA (blue) stained. Cells are viewed from two orientations as illustrated in the micrograph resulted from a 3-dimentional (3D) projection. The conspicuous appearance of MYA1 in the phragmoplast midzone is accompanied by the striking signal at the CDS (arrowheads) in a lateral view. In the axial view, the MYA1 signal forms discontinuous patches along the perimeter of the cell and such a pattern can also be detected in the 3D projection. (E) The MyoXI-4KO mutant forms misoriented cell plates when challenged by oryzalin. Root cells of wild-type (i), MyoXI-4KO (ii), and MyoXI-4KO expressing MYA1-GFP (iii) are outlined by propidium iodide staining. The mutant cells (ii), but not the control or rescued ones, produces extensive misoriented cell plates in cells marked by asterisks. The degree of misorientation is also reported quantitatively in Supplemental Figure 1B. (F) The MyoXI-4KO mutant loses the expansion guidance of the cell plate marked by GFP-SYP111 upon oryzalin treatment. A root cell forms a tilted SYP111-labeled cell plate (green) which seeks maximum expansion in the diagonal direction (arrows). Following the expression of the MYA1-TagRFP fusion protein (red), the expanding cell plate marked by GFP-SYP111 (green) aims to unify with the CDS marked by MYA1-TagRFP.

Oryzalin hypersensitivity could reflect cell division defects, which prompted us to examine whether MYA1 participates in cell division. In a transgenic line expressing MYA1-GFP and mCherry-TUB6 (β-tubulin 6), MYA1 exhibited a cell cycle-dependent pattern of dynamic localization corresponding to the reorganization of mitotic microtubule arrays (Figure 1B; Supplemental movie 1). At prophase when the mature PPB was present (asterisks, Figure 1B), MYA1-GFP did not exhibit a discernable localization pattern at a specific site. At later stages of prophase when the bipolar microtubule array establishes on the nuclear envelope and the PPB began to disassemble, MYA1-GFP became enriched in the cortical position occupied by the PPB (arrows, Figure 1B). This conspicuous localization of MYA1-GFP at the PPB site persisted throughout mitosis, until the expanding phragmoplast contacted it (arrowhead, Figure 1B). MYA1-GFP was also enriched in the midzone of the metaphase spindle and became obvious on microtubule bundles in the spindle midzone at anaphase (00:09:20 to 00:14:20, Figure 1B). The signal appeared more striking in the midzone of the developing phragmoplast and it expanded with the microtubule array towards the CDS (00:20:20, Figure 1B). This phragmoplast midzone-associated MYA1 signal eventually unified with that at the cortex (00:39:20, Figure 1B). Such a dynamic redistribution was further revealed by kymograph illustrating the appearance of the cortical signal (arrowheads) and the unification of the two spatially separated signals towards the end of cytokinesis (Figure 1C).

To determine how low doses of oryzalin compromised root growth in MyoXI-4KO mutant, we examined mitotic microtubule arrays by live-cell imaging after having a visGreen-TUB6 (β-tubulin 6) expressed (Supplemental Figure 2, Movie 2). Although the mutant cells did not exhibit noticeable differences in mitotic microtubule arrays when compared to the complemented cells expressing identical MYA1-GFP and mCherry-TUB6, they suffered severe challenges in establishing robust bipolar spindle microtubule arrays (−00:11:05 to 00:27:04, Supplemental Figure 2, Movie 3). While the PPB array was detected (arrowheads, −00:11:05, Supplemental Figure 2), very little if any microtubules were detected on the nuclear envelope at all stages of prophase. Then, conspicuous microtubules were assembled in the cell center and organized into flattened patterns that were hardly recognized as bipolar arrays (00:25:34-00:27:04, Supplemental Figure 2, Movie 4). The phragmoplast array formed and expanded but was turned 90°, perpendicular to the PPB-defined plane (00:34:35-00:45:52, Supplemental Figure 2). Although the mitotic cell in the rescued line showed a similar delay in completing cell division, its spindle microtubule array was able to recover and reestablish bipolarity, resembling those in cells treated with DMSO (Supplemental Figure 2, Movie5). Furthermore, the rescue cell showed no obvious defects in phragmoplast guidance. We conclude that the loss of the four Myosin XI motors caused serious defects in phragmoplast guidance as well as the assembly of bipolar spindle microtubule array that became vulnerable to subtle challenges on microtubules.

To capture the relationship between MYA1 and the developing phragmoplast microtubule array, we performed immunofluorescence experiments in isolated root cells to improve the signal-to-noise ratio. Since individual cells are isolated, they can be mounted on a microscope slide in any orientations. In a lateral view parallel to the cell division axis, MYA1 was detected at the CDS (arrowheads) besides being enriched in the phragmoplast midzone (Figure 1D). Cells that were placed with the cell division axis perpendicular to the glass slide permitted us to take an axial view across the cell cortex together with the phragmoplast midzone. In this informative view, MYA1 was detected as discrete patches forming an intermittent ring at the cell cortex (Figure 1D). These cortical patches are also observed in live cells, verifying our results from immunostaining is physiologically relevant (Supplemental Figure 3). In 3-D projections during cytokinesis, MYA1 was detected as a complete ring of patches along the CDS consolidated with the signal detected in the phragmoplast midline. (Figure 1D). Because Myosin XI-K was also detected at similar locations (*7*), this dual localization in the phragmoplast and at the CDS supported the notion that MYA1 together with other Myosin XI motors play a partially redundant role in the spatial regulation of cytokinesis.

We then tested whether the oryzalin-hypersensitive growth defects were brought about by cytokinetic defects. When root cells were outlined by the fluorescence dye propidium iodide, the oryzalin-treated MyoXI-4KO root cells often formed randomly oriented cell plates (asterisks, Figure 1E). Meanwhile, the wild-type control and the rescued line produced cells with uniform orientations in each cell layer under identical conditions (Figure 1E, Supplemental Figure 1B). Quantifications of cell wall angles showed that upon oryzalin treatment, MyoXI-4KO roots form misoriented cell walls at angles that varied from the typical 90° to the intercepting wall. To determine how the mutant cells produced such misaligned cell plates, we had the MyoXI-4KO mutant express a GFP fusion protein of the syntaxin protein SYP111/KNOLLE which served as a marker of the developing cell plate as reported (*15*). Upon oryzalin treatment, the MyoXI-4KO mutant cell formed the cell plate that sought for maximum expansion in a diagonal direction (arrows, Figure 1F; Supplemental movies 2). This cytokinetic defect was suppressed in mutants expressing a functional MYA1-TagRFP fusion protein which marked the CDS, as cells formed the regular transverse cell plate parallel to others in the same cell layer (arrowheads) (Supplemental Movie 3). This result suggested that the MyoXI-4KO mutant cells failed to form cell plates that are oriented perpendicular to the cell division axis. Furthermore, MYA1 marked precisely where the cell plate fused with the mother cell wall. In combination with MYA1 localization and the MyoXI-4KO cell plate misalignment, we concluded that MYA1 is a CDS protein that regulates the orientation of the cell division plane during cytokinesis.

Like MYA1, a few other proteins have been detected and persist at the CDS during mitosis in *A*. *thaliana*, including the microtubule-associated proteins MAP65-4 and TAN1 (Tangled 1) and the microtubule-based motor proteins POK1/2 (Phragmoplast Orienting Kinesin 1/2) (*16, 17*). Furthermore, Myosin XI-K also was detected at the CDS (*7*). To examine the relationship between MYA1 and these CDS-localized proteins, we performed dual-localization experiments in root meristematic cells from plants co-transformed with constructs expressing MYA1 and proteins of interest tagged with their respective fluorescent protein or the complementary 4xMyc tags. At the lateral view, the MYA1 signal overlapped with those of all four CDS proteins, Myosin XI-K, MAP65-4, TAN1, and POK1 (Figure 2A-D). Since these patches occupy the cortex as an intermittent ring, the axial view allowed us to examine these proteins across the entire perimeter at the cortex. All four CDS proteins formed discrete cortical patches aligned at the edge that colocalized with the MYA1 patches (Figure 2A-D). To further reveal the spatial relationship between MYA1 and these established CDS proteins in the axial view, fluorescence intensity scans of the CDS-localized signals were performed to graphically display the colocalizations in the form of overlapping peaks. It is worth noting that both MYA1, Myosin XI-K, and MAP65-4 were prominent at both the CDS and in the phragmoplast midzone, whereas TAN1 and POK1 were conspicuous at the CDS but not as noticeable in the phragmoplast. This phenomenon suggests that perhaps different assemblies of cytoskeletal factors are formed in the phragmoplast and at the CDS. Our results demonstrated the colocalization of MYA1 cortical patches with known CDS proteins. Therefore, we concluded that at least MYA1, XI-K, MAP65-4, TAN1, and POK1 assembled into discrete cytoskeletal patches at the CDS established by the PPB, which were termed as <u>C</u>ytoskeletal <u>M</u>otor <u>A</u>ssemblies at the <u>D</u>ivision <u>S</u>ite (CMADS), as a form of high order molecular assemblies.

**Figure 2.**
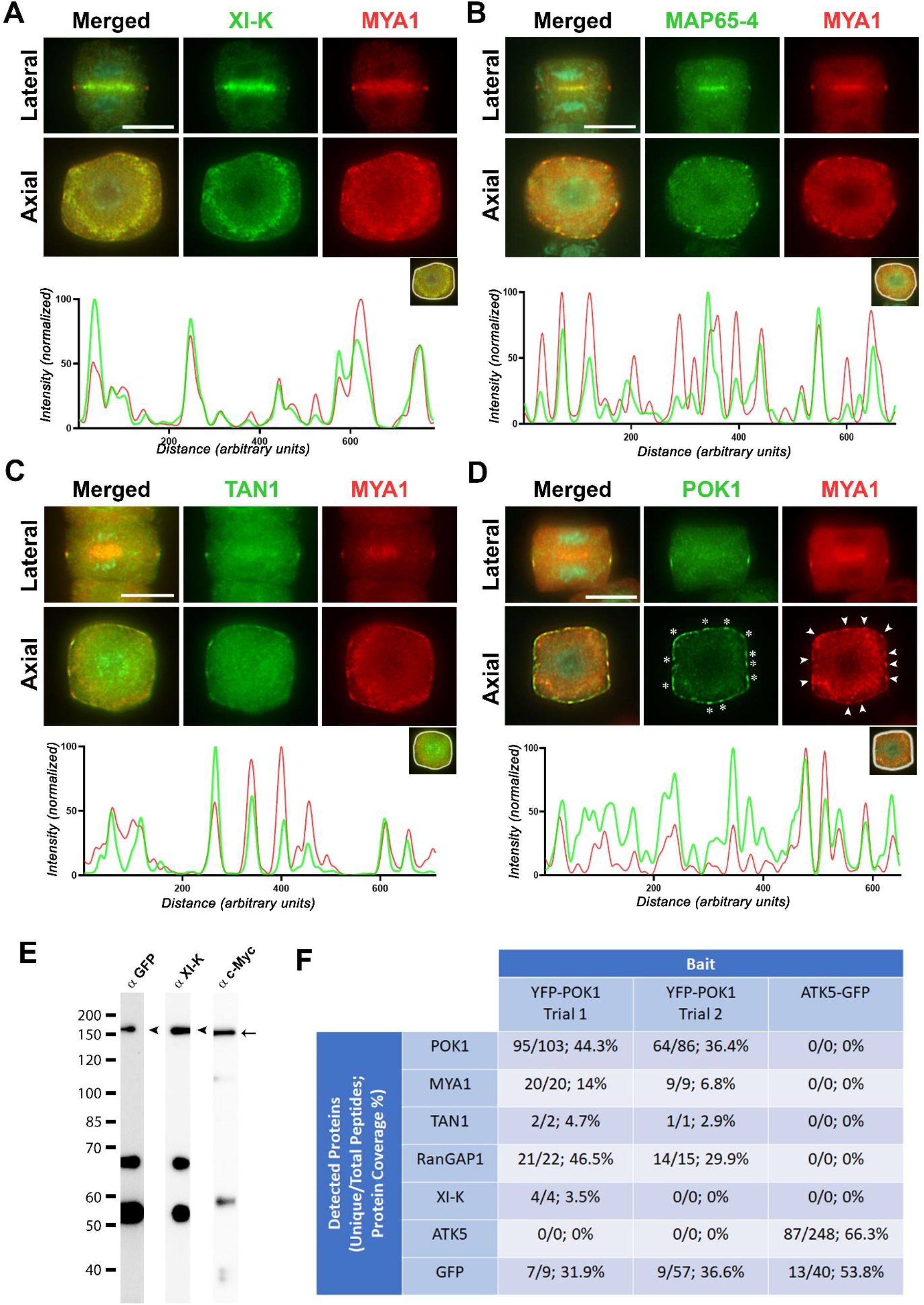
Myosin XI and other CDS cytoskeletal proteins form discrete patches of the Cytoskeletal Motor Assemblies at Division Sites (CMADS) *in vivo*. Scale bars: 5 μm. (A-D) MYA1 co-localize with other CDS proteins in cortical patches. Micrographs show dual localizations of MYA1-4xMyc (pseudo-colored in red) and XI-K-YFP (green) (A), MYA1-4xMyc (red) and MAP65-4-GFP (green) (B), MYA1-GFP (red) and TAN1-4xMyc (green) (C), and MYA1-4xMyc (red) and YFP-POK1 (green) (D) from lateral and axial views. The accompanying graphs on the bottom are results of fluorescence intensity scans of the two respective signals in the highlighted peripheral line encompassing the CMADS by using the merged images (inserts). The fluorescence intensities are normalized with the brightest signal set at 100 and the x axis has the cell perimeters set in arbitrary units. (E) MYA1 and Myosin XI-K associate with each other *in vivo*. Immunoprecipitation of Myosin XI-K-YFP, detected by both anti-GFP and anti-Myosin XI-K antibodies by immunoblotting (arrowheads), has MYA1-4xMyc co-purified and detected by anti-c-Myc antibodies (arrow). Note bands below 70-kDa are likely resulted from protein degradation. Both myosins are not detected when GFP is precipitated under similar conditions using transgenic plants expressing GFP alone (Supplemental Figure 3). (F) Immunoaffinity purification of YFP-POK1 reveals the association of MYA1, Myosin XI-K, POK1, TAN1, and RanGAP1 *in vivo*. When the YFP-POK1 fusion protein is purified, MYA1, TAN1, RanGAP1, and Myosin XI-K (XI-K) can be detected when assisted by LC-MS-MS analysis. Detected total and specific peptides as well as the coverage of polypeptide sequence in percentages are listed. A similar purification aimed at ATK5-GFP (*19*) is used as a reference for the determination of the specificity of detected proteins.

To test whether the Myosin XI motors described above physically associated with each other *in vivo*, we performed anti-GFP immunoprecipitation experiments using transgenic plants coexpressing Myosin XI-K-YFP and MYA1-4XMyc. The bait of XI-K-YFP fusion protein was detected by the anti-GFP antibodies as well as polyclonal antibodies raised against a Myosin XI-K specific peptide (arrowheads, Figure 2E). The precipitates also included the MYA1-4XMyc fusion protein which was revealed by a polyclonal anti-c-Myc antibody (arrow, Figure 2E). These myosin fusion proteins were specifically isolated because they were absent when similar anti-GFP immunoprecipitation experiments were performed by using extracts of plants expressing GFP alone (Supplemental Figure 4). Therefore, we conclude that the two actin motors associated with each other *in vivo*. The striking colocalization of MYA1 and the established CDS proteins inspired us to test whether it was due to physical association. To test this hypothesis, we performed an anti-GFP co-immunoprecipitation experiment using plants expressing YFP-POK1 (*17*). POK1 was chosen due to its elevated expression in mitotic cells as well as being primarily localized at the CDS, making it the candidate of choice to dissect protein association in CMADS. We were able to detect by mass spectrometry the bait protein POK1 as well as the YFP/GFP tag with high coverages of the polypeptides (Figure 2C). When the co-purified proteins were subject to peptide identification, MYA1 was recovered with 20 unique peptides detected (Figure 2F). Together, Myosin XI-K as well as the known POK1-interacting TAN1 were also detected. We also detected the RanGAP1 protein with greater than 46% coverage of the polypeptide. RanGAP1 was previously detected at the CDS and physically interacts with the C-terminal coiled-coil domains of POK1 (*18*). Therefore, RanGAP1 is hypothesized to be a bona fide component of the CMADS. Results obtained from the purification of mitotic kinesin motor ATK5 served a negative control because it is associated with the spindle apparatus but not detected in the spindle midzone or the CDS (*19*). The cross-examination verified that the proteins detected at the CMADS were specifically co-purified with POK1 as none were detected in when ATK5-GFP was purified under identical conditions (Figure 2F). Together with co-localization evidence, our results here further support the notion that the CMADS harbors, but is not limited to, Myosin XI and Kinesin-12 motor together with other cytoskeleton-associated factors like TAN1 and RanGAP1.

Because MYA1 localized to the PPB site and consequently was detected at the CDS at later stages of mitotic division, we asked how the disturbance of PPB assembly would affect MYA1 localization. Three functionally redundant microtubule-associated proteins TRM6, TRM7, and TRM8 (TON1 Recruiting Motif 6, 7, 8) play critical roles in the assembly of the PPB microtubule array as the *trm678* triple mutant exhibits various degrees of PPB defects (*20*). We examined MYA1 localization in the *trm678* triple mutant cells and found that the MYA1 signal was still detectable at the CDS although it often appeared at one side only in a lateral view of the dividing cells while its localization in the phragmoplast midzone was still conspicuous (Figure 3A). Such a phenomenon prompted us to examine cells from an axial view. We found isolated, sparse MYA1 patches in the cell perimeter representing the CDS at the focal plane (Figure 3A). To verify that the abnormal distribution of MYA1 patches was caused by the loss of the TRM function, we employed a complemented line in which TRM7-3xYFP and MYA1-4xMyc were expressed in the *trm678* triple mutant. TRM7 is not detectable at the CDS at anaphase (*20*), therefore, we examined metaphase cells from both lateral and axial views. In this complemented line, TRM7 was detected at the CDS like MYA1 in the lateral view (Supplemental Figure 5). The axial view shows that remnant TRM7 signal was detected across the CDS while the MYA1 cortical patches were clearly formed by metaphase with near regular distances across the cell perimeter (Supplement Figure 5). As a control for our quantifications, we examined MYA1 patches in our MYA1-GFP rescue line, which developed normal PPB (asterisks, Supplement Figure 2). To report the difference in MYA1 localization in the *trm678* mutant and rescue line, we quantified the number of MYA1 patches per 10 μm of the cell perimeter. The number dropped significantly to an average of fewer than 2 (1.573 ± 0.161) patches/10 μm in the *trm678* mutant while the rescue cell formed greater than 3 (3.614 ± 0.186) patches/10 μm (Figure 3B). Because the arrangement of the MYA1 patches became irregular in the *trm678* mutant, the distances between adjacent patches were measured by using the cell perimeter as a reference.

**Figure 3.**
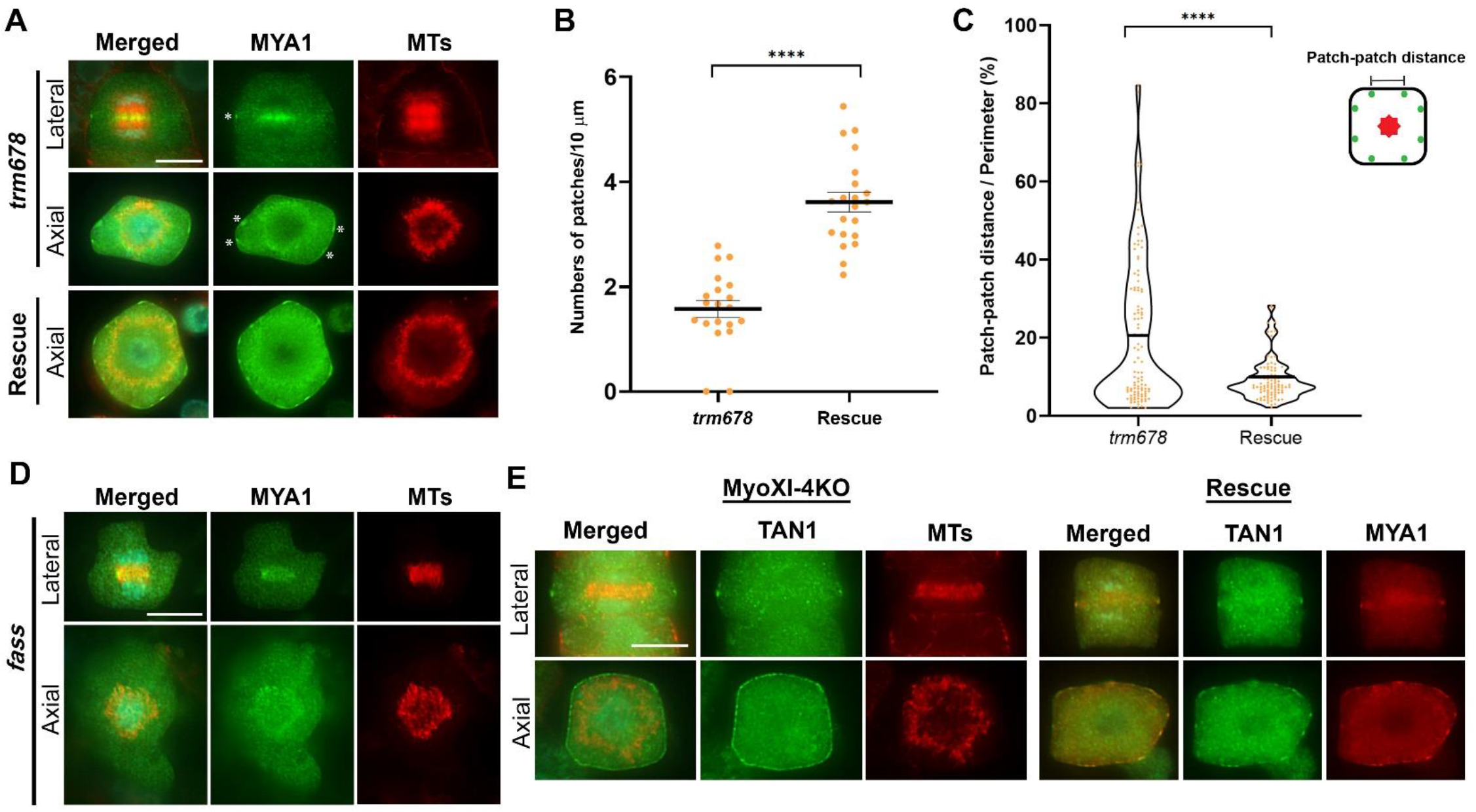
MYA1 localization and CMADS formation at the cortex are dependent on the PPB. Scale bars: 5 μm. (A) MYA1 localization at the CDS is disrupted in the *trm678* mutant cells. When examined from the lateral view, the MYA1 (green) signal often stands out only at one side of the mutant cell (asterisk) while its presence in the midline of phragmoplast microtubule (MTs) array (red) remains conspicuous. An axial view shows sparse, irregularly distanced MYA1 patches at the cell cortex (asterisks). The pattern of regularly distanced MYA1 patches is restored in cells expressing TRM proteins (rescue). (B) The numbers of patches are determined in both the *trm678* mutant and the rescue line and are presented quantitatively per 10 μm. Sample sizes have n ≥ 20 cells per genotype. The difference is determined significant based on a two-tailed t-test with a P-value < 0.0001 (****) (C) The patterns of MYA1 patches in the *trm678* mutant and rescue line are assessed in the form of patch to patch distance when the distances between greater than 90 adjacent patches were measured in more than 10 cells of each genotype. The difference is determined significant based on a two-tailed t-test with a P-value < 0.0001 (****). Means are marked by solid black line. (D) MYA1 no longer form cortical patches in the *fass* mutant lacking the PPB. While MYA1 (green) can be detected in the association with phragmoplast microtubules (red) in both lateral and axial views, no discernable signal stands above the background at the cell cortex. (E) TAN1 localization at the cell cortex is altered in the MyoXI-4KO mutant. Although TAN1-4xMyc (green) is detected at the CDS as shown in the lateral view, it becomes nearly continuous across the CDS from the axial view. In the cell of the rescued plant (rescue)expressing MYA1-GFP (red), the discrete TAN1-4xMyc (green), however, discrete patches are detected in the axial view. Scale bar: 5 μm

While the rescue cells had distances approximately 9% of the perimeter (9.256% ± 0.554 %), the *trm678* mutant cells had the distances more than doubled and varied significantly averaging at approximately 20% of the perimeter (20.201% ± 1.83%) (Figure 3C). Therefore, the results suggested that defects of PPB assembly affected the formation of the otherwise regularly spaced MYA1 patches at the CDS in the *trm678* mutant. To take one step further, we then asked whether the PPB was essential for MYA1 to assume the localization at the CDS by employing the *fass* mutant which fails to establish the PPB at the cell cortex (*21*). When MYA1-GFP was expressed in the *fass* mutant cells, it was clearly detected in the phragmoplast but became undetectable at the cell cortex in both lateral and axial views (Figure 3D). These results led to the conclusion that the PPB is required for MYA1 localization at the CDS. Conversely, we asked if the formation of the CMADS might be affected in the MyoXI-4KO background. TAN1 was used as a marker of CMADS and was still detected at the CDS in both lateral and axial views in the MyoXI-4KO mutant (Figure 3E). However, the discrete TAN1 patches were replaced by nearly continuous signal across the perimeter in the MyoXI-4KO mutant cells. The rescued plants expressing a functional MYA1 fusion protein had the TAN1 signal return to discrete patches (Figure 3E). Taken together, these results led to the conclusion that the PPB was required for the initial localization of CMADS to the CDS, and Myosin XI played a role in organizing these CDS-localized proteins into discrete CMADS appearing in patches.

F-actin exhibits a dynamic reorganization pattern at the cell cortex during mitosis, first colocalizes with the PPB and later disappears from the mature PPB and remains largely absent at the PPB-defined CDS for much of the later mitotic stages (*22*). Because F-actin functions as myosin tracks and plays essential role in cell division plane orientation, we asked whether the CMADS formation depended on F-actin. When F-actin was depolymerized by latrunculin B (Lat B), the MYA1 signal was dispersed vertically in the cell division axis at the cell cortex, when compared to consolidated MYA1 signal at the CDS in the mock/DMSO-treated cells (Figure 4A). To examine whether the signal diffusion only took place along the division axis, the cells were also examined from an axial view (Figure 4A). We found that the discrete MYA1 patches were replaced by nearly continuous signal across the cell perimeter after Lat B treatment, instead of being in discrete patches (Figure 4A). Intensity scan of the MYA1 fluorescent signal across the perimeter in the axial view demonstrated the peaks and troughs reflecting the patches formed in the DMSO-treated cell, whereas the Lat B-treated cell displayed a close to linear intensity pattern due to the diffused signal (Figure 4B). To further assess the vertical diffusion of the MYA1 signal, we measured the width of the MYA1 signal in the lateral view by using the total perimeter of the cell in this same focal plane as a reference. In the Lat B-treated cells, the MYA1 signal diffused more than two-folds by occupying 4.5% of the perimeter (4.583% ± 0.2578%), compared to having the signal confined within approximately 2% of the cell perimeter (2.02% ± 0.087%) in the mock-treated cells (Figure 4C). Because CMADS consist of different cytoskeletal proteins, we asked if POK1 also required F-actin to be concentrated in discrete cortical patches by co-expressing both YFP-POK1 and MYA1-4xMyc. Like MYA1, the CDS-localized POK1 requires F-actin to become concentrated in the lateral view and consolidated into discrete patches in the axial view (Supplemental Figure 6). The results allowed us to conclude that intact F-actin filaments were essential for the consolidation of CMADS both lateral and axial axes. Finally, we asked whether the actin-dependent CMADS formation was coupled with division plane determination by employing cells expressing both MYA1-GFP and mCherry-TUB6 to monitor myosin localization with microtubule arrays as references. After Lat B treatment, MYA1 was detected in the spindle midzone but not at the cell cortex prior to phragmoplast assembly (−00:07:39 to 00:09:55, Figure 4D). Following mitosis, the developing phragmoplast became tilted while the diffuse MYA1 signal was detected at the cell cortex (brackets, 00:25:31, Figure 4D). Towards the end of cytokinesis, the phragmoplast microtubule array and the associated MYA1 signal remained tilted, irrespective of the orientation of the CDS-localized MYA1 signal (00:38:17, Figure 4D). In the mock-treated cells, in contrast, the MYA1-GFP signal became concentrated at the CDS around the time of nuclear envelope breakdown (00:00:00, arrowheads, Figure 4D). The CDS-localized MYA1 signal persisted at later stages of mitosis and became most conspicuous around the time when the phragmoplast microtubule array was expanding centrifugally (asterisks, 00:15:20, Figure 4D). Concomitantly with the expanding microtubule array, the phragmoplast-localized MYA1 signal was always aligned with the MYA1-marked CDS, and eventually merged with the cortical signal (00:30:25, Figure 4D). We concluded that the diffuse cortical MYA1 signal caused by Lat B was no longer recognized by the developing phragmoplast.

**Figure 4.**
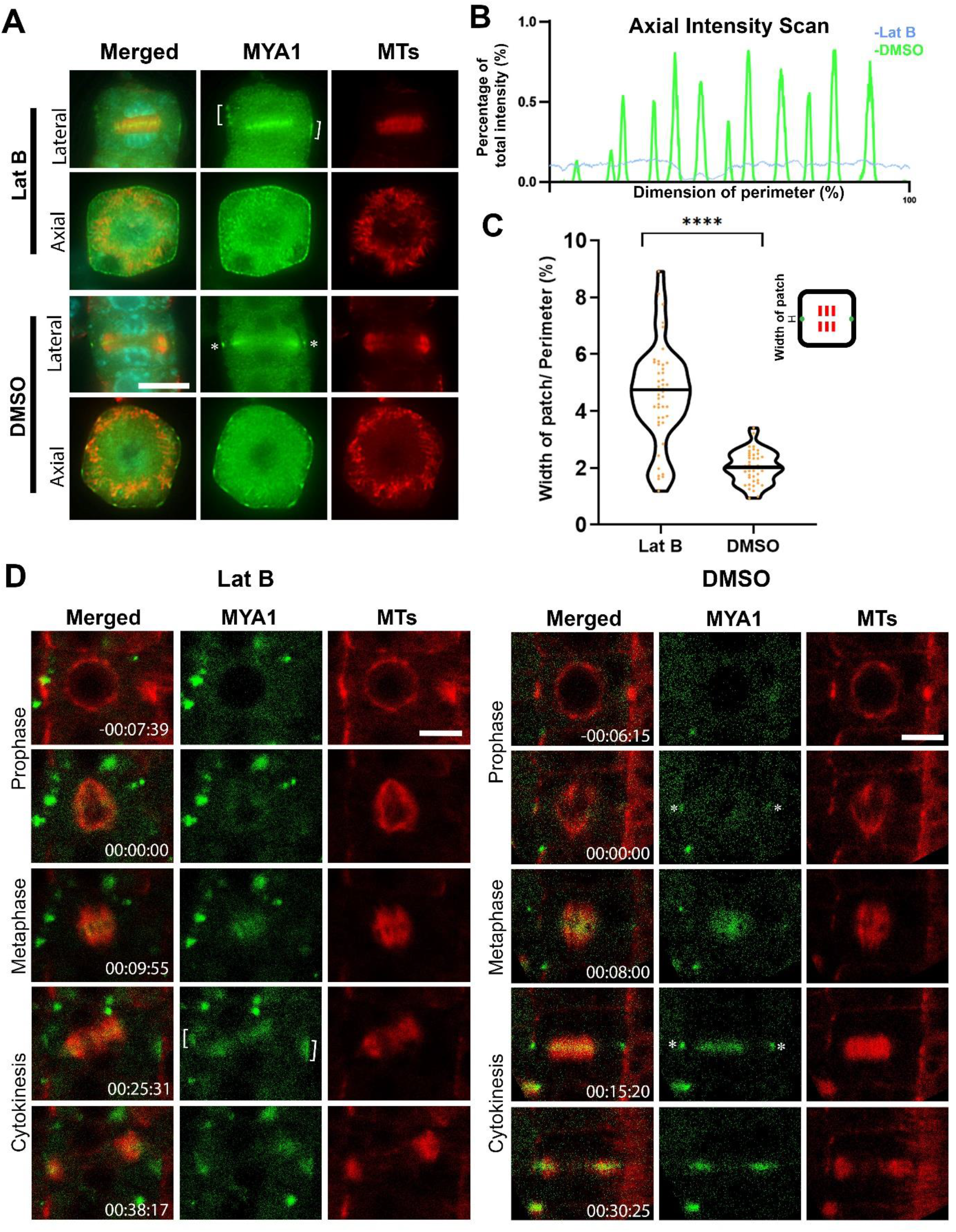
CMADS consolidation requires intact F-actin. All micrographs have MYA1 pseudo-colored in green and microtubules in red. Scale bars: 5 μm. (A) Latrunculin B (Lat B) treatment causes diffusion of the MYA1 signal at the CDS. MYA1 appears in expanded zones (brackets) following the treatment when the cell is examined from a lateral view, although its appearance in the phragmoplast midzone remains concentrated. From an axial view, MYA1 loses its appearance in discrete patches which are replaced by nearly continuous signal across the perimeter. Mock-treated cells exhibit normal localization pattern of MYA1 in consolidated patches at the CDS. The micrographs have MYA1 pseudo-colored in green and microtubules (MTs) in red, and DNA in cyan in the merged images. (B) Assessment of MYA1-GFP signal distribution across the cell perimeter. The fluorescent signal is reported as the percentage of the sum of the MYA1-GFP signal. The x axis represents the dimension of the cell perimeter from point 0 to 100% of the perimeter. (C) Quantitative assessment of the width of the MYA1 patches shows significant expansion along the division axis. The occupation of the cell perimeter by the width a MYA1 patch is determined in the percentage in the lateral view. Each treatment has the sample size n > 40 cells. The difference in MYA1 signal occupancy of the cell perimeter (in percentage) between mock and Lat B treated cells is determined significant by using a two-tailed t-test with a P-value < 0.0001 (****). Means are marked by solid black line. (D) Lat B treatment causes diffusion of MYA1 at the CDS in transgenic cells expressing both MYA1-GFP and mCherry-TUB6. In a mitotic cell treated with Lat B, the MYA1 signal is readily detected at the cell cortex in a diffuse form (brackets) when the phragmoplast is formed. In mock (DMSO)-treated cell, MYA1 is detected at the CDS as consolidated signals (asterisks). In the meantime, MYA1 localization in the phragmoplasts after both Lat B and DMSO treatments do not show noticeable differences by live-cell imaging. The colored images have MYA1-GFP pseudo-colored in green, microtubules (MTs) in red.

Our results summarized above revealed the dynamic process of high order cytoskeletal assemblies of both microtubule- and F-actin-associated factors at the CDS and both the localization of these factors and the formation of this CMADS were brought about by the PPB (Supplemental Figure 7). Early, broad PPB microtubule arrays recruit microtubule-associated proteins like MAP65-4 to bridge neighboring microtubules and TAN1 to attract the kinesin POK1. Following the condensation of the PPB microtubule array, probably brought about by MAP65-4 *via* its microtubule-bundling activities along with F-actin function (*9*), brings in additional cytokinesis important factors like RanGAP1 and Myosin XI motors (Supplemental Figure 7). We also demonstrated that intact F-actin and Myosin XI motors are essential for the CMADS to become concentrated at spatially confined sites at the CDS and detected a novel function of Myosin XI motors in the robust reorganization of the spindle MT array. The spatial consolidation of the CMADS was required for the recognition of the CDS by the expanding phragmoplast during cytokinesis. Our discoveries led to the conclusion that the CMADS translate the positional information of the transient PPB microtubule array into cell plate fusion location at the CDS, an essential action for the determination of the cell division plane during cytokinesis. Furthermore, MYA1, as an F-actin-based motor that colocalized with microtubule arrays, serves as an excellent candidate that establishes the crosstalk between two dynamic cytoskeletal filaments at the CDS as well as in the phragmoplast midzone during cytokinesis. This study, therefore, inspires us to pursue what the complete constituents of the CMADS are and how proteins of different cytoskeletal elements congregate to form the dynamic CMADS for spatial regulation of cytokinesis in land plants.

## ACKNOWLEDGEMENTS

This work was supported by the NSF grant MCB-1920358 and the U. S. Department of Agriculture (USDA)-the National Institute of Food and Agriculture (NIFA) under an Agricultural Experiment Station (AES) hatch project (CA-D-PLB-2536-H). We thank Dr. John Harada for critical comments. We are indebted to Dr. Valerian V. Dolja for sharing the Myosin XI constructs and seeds, Dr. David Bouchez and Dr. Martine Pastuglia for the *trm*, *ton* and *fass* lines, Dr. Sabine Müller for the YFP-POK1 line, Dr. Masa H. Sato for the GFP-SYP111 plasmid, and Dr. Tsuyoshi Nakagawa for the pGWB plasmids. Supplemental Figure 7 was made by using the BioRender.com tool.

## Notes

### Competing Interest Statement

The authors have declared no competing interest.

