## Supplemental Figures for "The microtubular preprophase band recruits Myosin XI to the division site for plant cytokinesis"

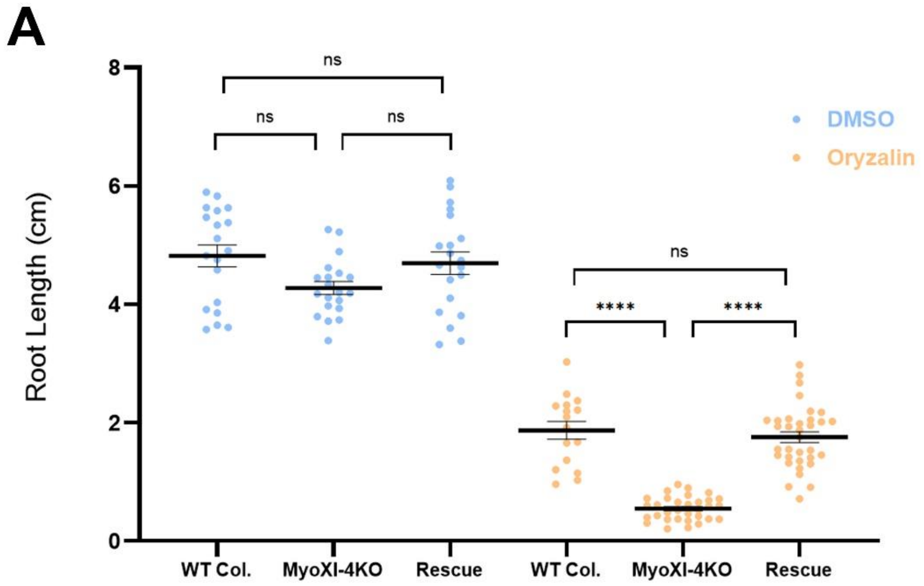


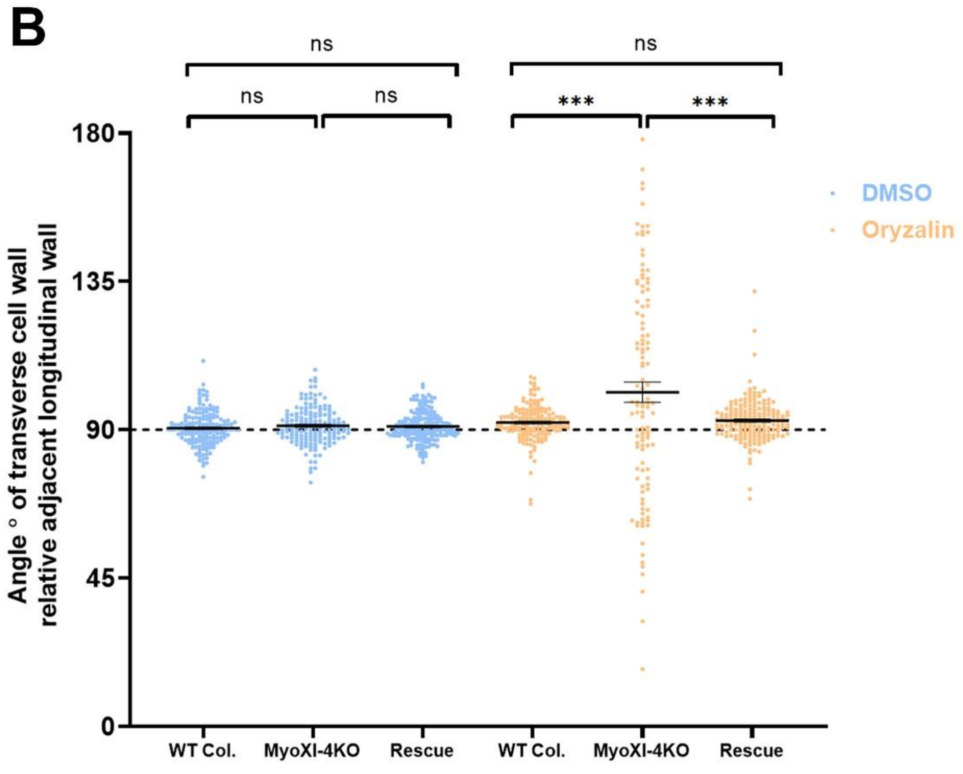

**Supplemental Figure 1.** Quantitative assessment of compromised root growth and altered division plane orientation in the MyoXI-4KO mutant.

(A) Root lengths in cm are measured in wild-type (WT Col), MyoXI-4KO, and MyoXI-4KO expressing MYA1-GFP (rescue). The MyoXI-4KO mutant produced roots slightly shorter than WT and the rescue line and the root extension phenotype is significantly enhanced after oryzalin treatment. Sample size n>15 per genotype. Differences between root length was determined by using a two-tailed t-test: P-value < 0.0001 (****), or no significance (ns).

(B) Orientation of the cell plate is assessed by measuring the angle to the adjacent cell wall. The MyoXI-4KO mutant root produces cell plates with great angle variations while the wild-type and rescue lines produce angles of close to 90°. Sample size n≥115 angles from 5 plants of each genotype. Differences between cell wall angle was determined by using a two-tailed t-test: P-value < 0.001 (***) or no significance (ns).

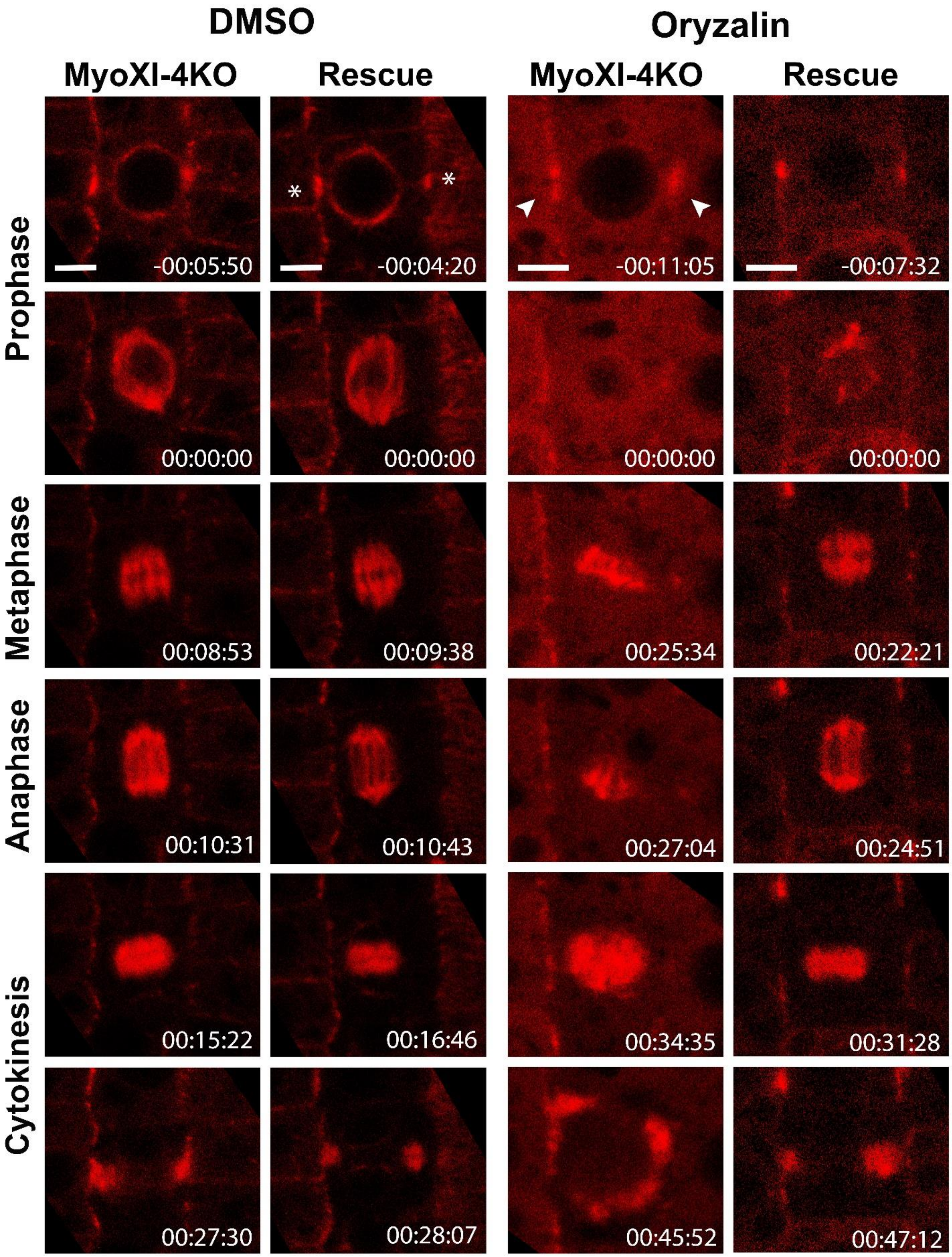
DMSO
Oryzalin
MyoXI-4KO
Rescue
MyoXI-4KO
Rescue
Prophase
Metaphase
Anaphase
Cytokinesis
-00:05:50
-00:04:20
-00:11:05
-00:07:32
00:00:00
00:00:00
00:00:00
00:00:00
00:08:53
00:09:38
00:25:34
00:22:21
00:10:31
00:10:43
00:27:04
00:24:51
00:15:22
00:16:46
00:34:35
00:31:28
00:27:30
00:28:07
00:45:52
00:47:12

**Supplemental Figure 2.** Myosin XI plays a role in reorganization of the spindle microtubule array during mitosis. The MyoXI-4KO mutant and MyoXI-4KO expressing MYA1-GFP (rescue) lines express a visGreen-TUB6 or mCherry-TUB6 fusion protein that marks microtubule arrays in live-cells undergoing mitosis. Snapshots are shown with time stamps in hr:min:sec. The MyoXI-4KO cell forms disorganized, compressed microtubule arrays during mitosis after oryzalin treatment while the cell of rescue line assembles bipolar spindle microtubule arrays. The oryzalin-treated mutant cell but not that of the rescue line forms a flipped phragmoplast microtubule array during cytokinesis. Such a difference is not obvious in cells treated with DMSO. Scale bar: 5 µm.

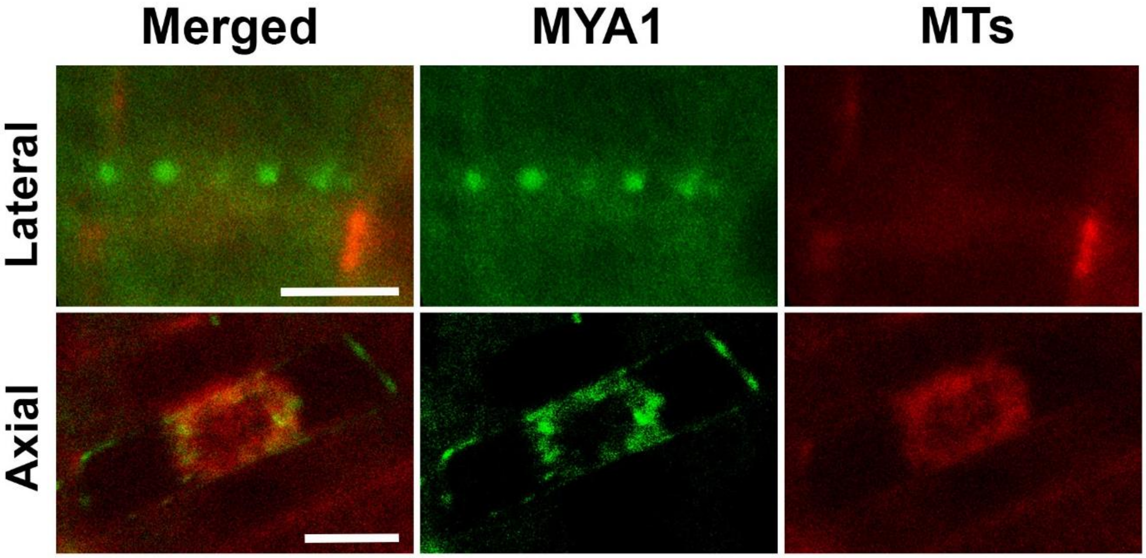


**Supplemental Figure 3.** The fluorescent signal of MYA1-GFP is concentrated in patches at the cortical division site (CDS) in living cells co-expressing mCherry-TUB6, with MYA1 pseudo-colored in green and MTs in red. In the lateral view, patches of MYA1-GFP signal are detected at the CDS where no obvious microtubules are detected. In the axial view, MYA1-GFP shows dual localizations of patches at the CDS and the signal associated with the phragmoplast microtubule array in the interior part of the cell. Scale bar: 5 µm.

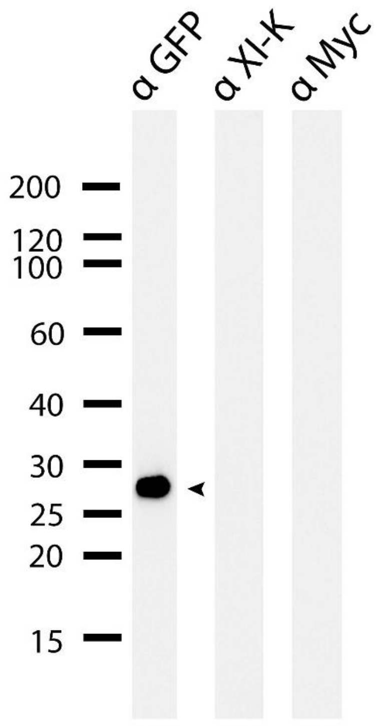


**Supplemental Figure 4.** Negative control of anti-Myosin XI-K immunoblotting. Free GFP protein is detected by anti-GFP immunoblotting after its precipitation from extracts of transgenic plants expressing the fluorescent protein alone.

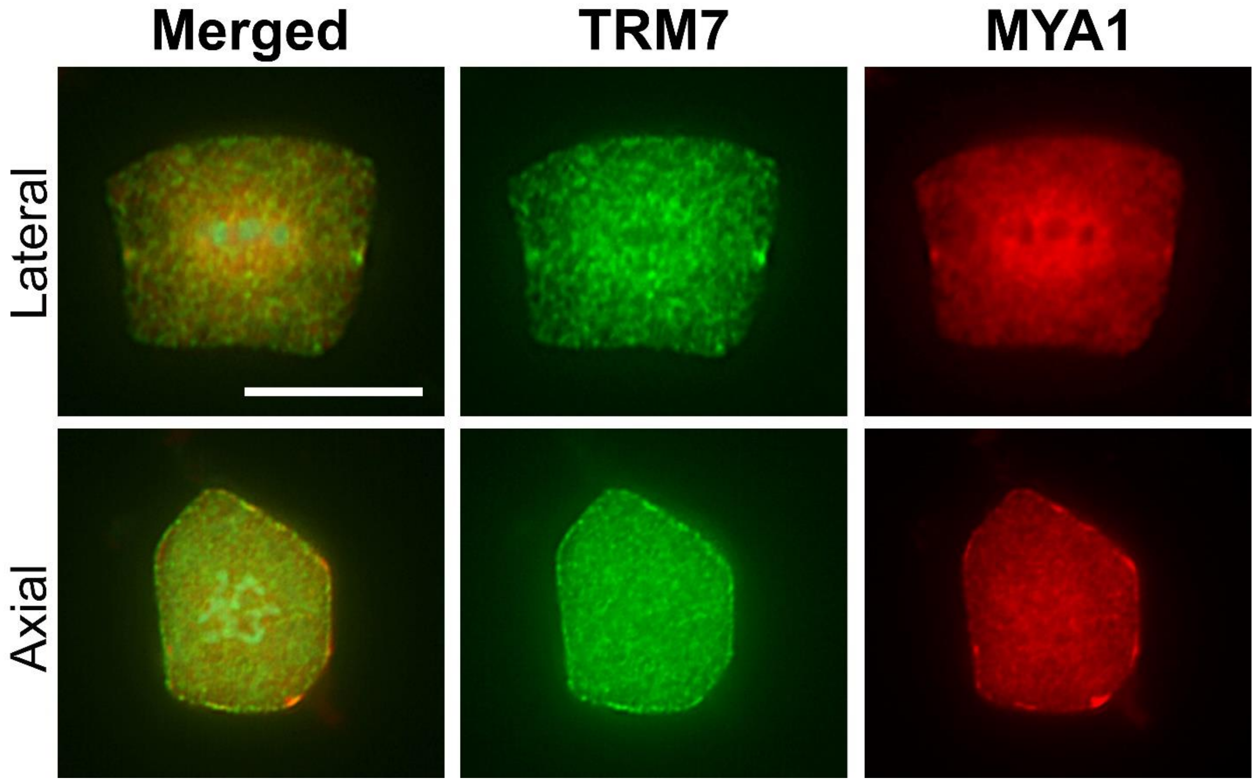


**Supplemental Figure 5.** Co-localization of TRM7 and MYA1 in metaphase cells. The TRM7-3xYFP fusion protein (green) is detected at the CDS where MYA1-4xMyc (red) is also found from the lateral view. While MYA1 is detected in patches from the axial view, remnants of TRM7 signal is detected at the cell cortex. Scale bar: 5 µm.

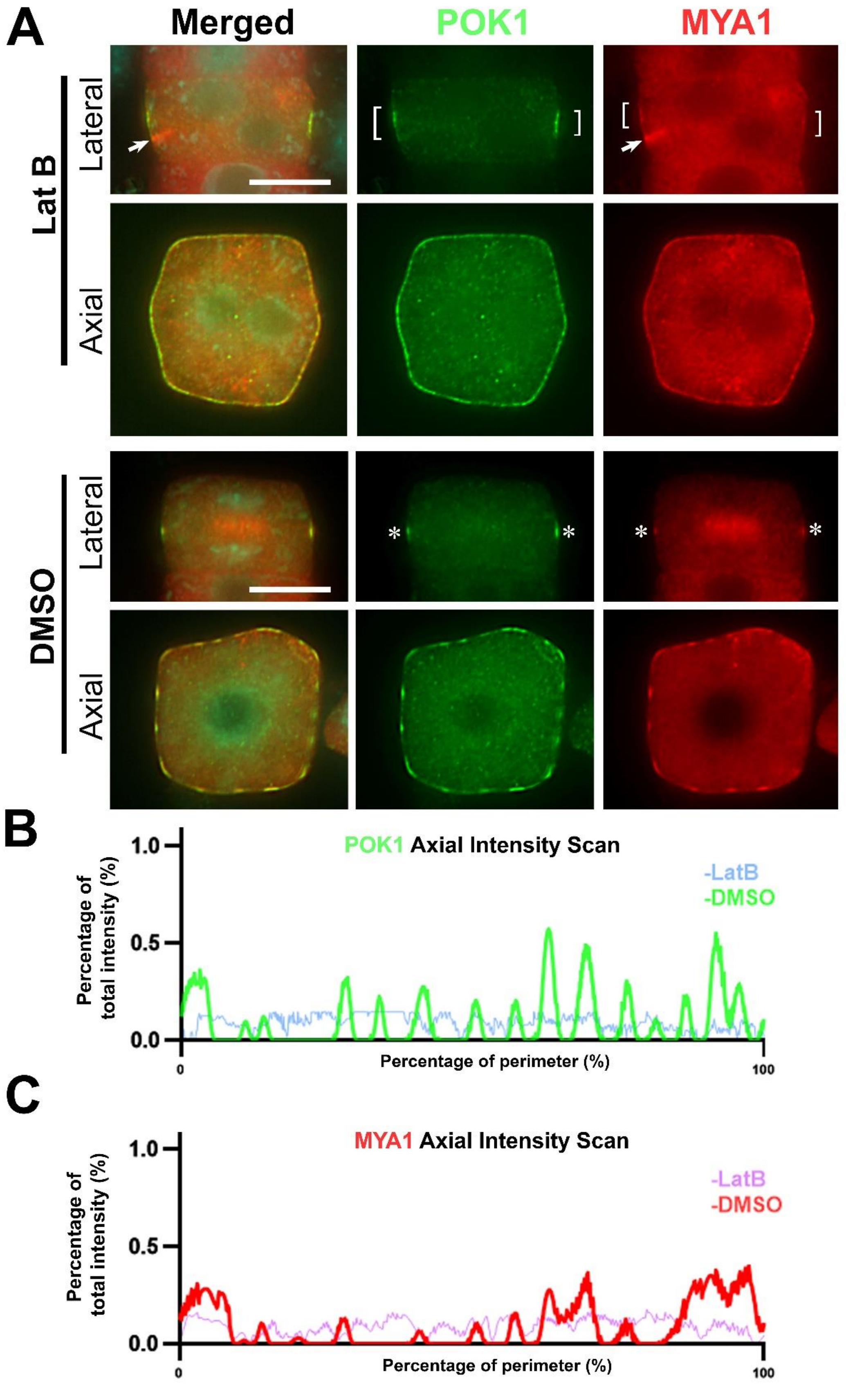

A
Merged
POK1
MYA1
Lat B
Lateral
Axial
DMSO
Lateral
Axial
B
POK1 Axial Intensity Scan
-LatB
-DMSO
Percentage of total intensity (%)
1.0
0.5
0.0
0
Percentage of perimeter (%)
100
C
MYA1 Axial Intensity Scan
-LatB
-DMSO
Percentage of total intensity (%)
1.0
0.5
0.0
0
Percentage of perimeter (%)
100

**Supplemental Figure 6.** Depolymerization of F-actin leads to diffusion of POK1 signal at the CDS. (A) Cytokinetic cells are examined from both lateral and axial views after the YFP-POK1 (green) and MYA1-4-cMyc (red) fusion proteins are detected by immunofluorescence. The merged images have DAPI-stained DNA in cyan. After Lat B treatment, POK1, like MYA1, is detected in widened zones (brackets) along the cell division axis. The phragmoplast midzone-localized MYA1 (arrows) misses the widened cortical POK1/MYA1-defined zone. In the axial view, POK1 and MYA1 localizes across the cell perimeter diffusely. In contrast, mock/DMSO-treated cells, POK1 and MYA1 mark the CDS (asterisks) from the lateral view and their signals are consolidated in patches across the cell perimeter in the axial view. (B, C) Assessment of the distributions of the POK1 (B) and MYA1 (C) signals across the cell perimeter in Lat B or DMSO-treated cells. The fluorescent signal is reported as the percentage of the sum of the POK1 and MYA1 signals, respectively. The x axis represents the dimension of the cell perimeter from point 0 to 100% of the perimeter. Scale bars: 5 µm.

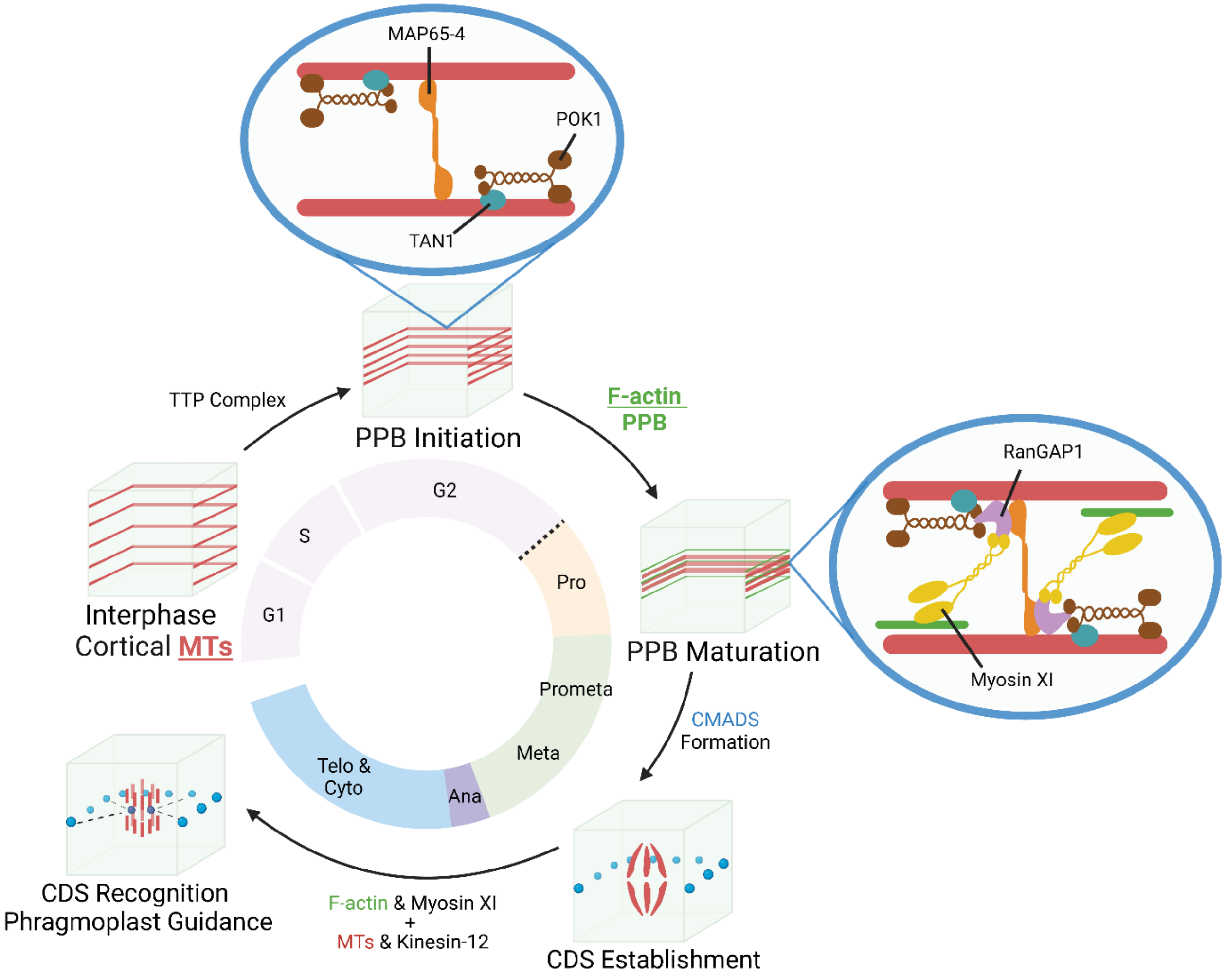


**Supplemental Figure 7.** Schematic presentation of CMADS formation and phragmoplast guidance during mitotic division in somatic plant cells. Following the TTP (TON1-TRM-PP2A) complex-dependent induction of PPB formation, microtubule associated factors like MAP65-4 and TAN1 together with its associated POK1 kinesin are recruited to the wide PPB. Narrow PPB attracts RanGAP1 and its associated Myosin XI to existing microtubule-interacting factors. F-actin in the PPB allows Myosin XI to consolidate these cytoskeletal factors into CMADS. CMADS persist at the CDS at later stages of mitosis until the phragmoplast midzone-localized MYA1 and other proteins unify with those in the CMADS in order to allow the expanding cell plate to insert into this PPB-defined site towards the end of cytokinesis.
